# Knowledge Graph-based Recommendation Framework Identifies Novel Drivers of Resistance in EGFR mutant Non-small Cell Lung Cancer

**DOI:** 10.1101/2021.07.23.453506

**Authors:** Anna Gogleva, Dimitris Polychronopoulos, Matthias Pfeifer, Vladimir Poroshin, Michaël Ughetto, Benjamin Sidders, Jonathan R. Dry, Miika Ahdesmäki, Ultan McDermott, Eliseo Papa, Krishna Bulusu

## Abstract

Resistance to EGFR inhibitors (EGFRi) presents a major obstacle in treating non-small cell lung cancer (NSCLC). One of the most exciting new ways to find potential resistance markers involves running functional genetic screens, such as CRISPR, followed by manual triage of significantly enriched genes. This triage process to identify ‘high value’ hits resulting from the CRISPR screen involves significant manual curation that requires specialized knowledge and can take even experts several months to comprehensively complete.

To find key drivers of resistance faster we built a hybrid recommendation system on top of a heterogeneous biomedical knowledge graph integrating preclinical, clinical, and literature evidence. Genes were ranked based on trade-offs between diverse types of evidence linking them to potential mechanisms of EGFRi resistance. This unbiased approach identified 36 resistance markers from >3,000 genes, reducing hit identification time from months to minutes. In addition to reproducing known resistance markers, our method identified novel resistance mechanisms that we prospectively validated.

## Introduction

In this study we explore how a biological question can be translated into a recommendation problem. Traditionally recommendation systems have been used to help users discover relevant items among an overwhelming number of options [1]. By interacting with recommendations users provide either implicit or explicit feedback, which recommendation models use to personalise and improve predictions [2]. Information overload is particularly common in e-commerce [3], streaming [4] and social media applications [5], hence recommendation systems play key role in these industries. The biomedical domain on the other hand, is not seen as a typical application area for recommendation systems. Their usage so far is limited to a handful of recent studies: Ozsoy et al, applied collaborative filtering for drug repositioning problem [6]; Frainay et al, developed a network-based recommendation solution to enrich and interpret metabolomicc data [7]; Suphavilai et al, built a matrix factorization-based recommendation system to predict response of cancer drugs [8]; Radivojevic et al, developed an automated recommendation solution for synthetic biology [9]. The success of these pilot studies suggest there are potential opportunities for recommendation systems in the biomedical domain; the amount of biomedical data is growing exponentially and scientists could benefit from recommendation solutions that help them to navigate the data and reason about it.

Naturally, direct transfer of classic recommendation approaches to the biomedical domain is not trivial. Specifics of the problem space impose numerous challenges for a recommendation system practitioner, to name a few:

- an elementary unit of recommendation is not a simple self-contained item (e.g a gene), but rather a research direction accompanied by a biologically sound hypothesis;
- ultimate validation of recommendations is complex and often requires expensive and time-consuming laboratory experiments, as opposed to users just ’selecting’ an item in a common non-biological recommendation scenario;
- unlike traditional applications, in a biomedical setting both implicit and explicit feedback is scarce, making it harder to tune and train models;
- ground truths are scarce and in most cases context-dependent which renders training challenging;
- due to the high cost associated with accepting a recommendation, an increased emphasis is placed on explainability and exposing causal reasoning paths behind a recommendation.

Despite these challenges, wider adoption of recommendation approaches holds plenty of opportunities to support and accelerate biological research. To illustrate this point, in this study we focused on the problem of drug resistance in lung cancer. Our goal was to build a recommendation solution that finds key genes driving drug resistance.

Drug resistance is a complex biological phenomenon that hinders development of efficient and lasting cancer treatments [10]. Tumours recruit diverse strategies to escape selective pressure induced by drugs, such as changes in drug metabolism [11], inhibition of cell death [12], epigenetic alterations [13] or acquired mutations in drug targets [14]. Enhanced DNA repair and increased amplification of tumour driver genes also contribute to secondary resistance [15]. Genetic and epidemiological diversity of patients [16] further complicates the resistance landscape.

In this study we focus on non-small cell lung cancer (NSCLC) carrying activating mutations of the epidermal growth factor receptor (EGFR). It accounts for 15-20% of lung cancer patients [17]. Treatment with first or second generation EGFR tyrosine kinase inhibitors such as gefitinib, erlotinib or afatinib results in impressive response rates in patients initially [18], however, tumours quickly develop resistance to treatment. The majority of resistant cases are driven by accumulation of secondary mutations of EGFR gene, such as T790M, that prevent binding of EGFR TKI compounds [19]. Development of osimertinib, a third generation EGFR TKI, provided the ability to target such secondary EGFR mutations [20]. In fact, treatment with osimertinib significantly improved patient survival in first- or second-line therapy setting [21, 22]. However, therapy resistance prevails. Acquired mutations of EGFR such as C797S drive osimertinib resistance in 6-26 % of cases. Bypass pathway activation, amplifications of MET or mutations in PIK3CA have also been shown to contribute to resistance [23]. Still, in half of the cases the molecular resistance mechanisms remain unknown and promising markers could reside in a so called ’dark matter’ of the human genome [24].

A common strategy to find key drivers of acquired resistance is based on functional genomic screens, such as CRISPR screens [25]. CRISPR-Cas9 genome-wide knock out, knock down and knock-in screens have recently emerged as an efficient high-throughput technology to systematically investigate resistance mechanisms [25]. CRISPR screens can be applied in two ways to understand drug response and drug resistance. First, they can be used to identify alterations in genes that increase sensitivity of a cell to drug treatment. Here, researchers measure negative selection of modified genes in drug treatment. This approach can help to define therapeutic combinations that might increase response to treatment. Second, CRISPR screens are used to identify genes that drive drug resistance if altered. In this case the experimental setup mimics treatment scenarios in the clinic. In this approach, outgrowth or positive selection of drug resistant cells is measured and used to define mechanisms that drive resistance. These can be potentially targeted once resistance is established.

In these settings, a typical output of a CRISPR screen may identify many hundreds of resistance genes. To narrow down the list to the most promising, biologically plausible and actionable resistance genes, researchers have to perform manual triage and validation. During this process experts aggregate prior knowledge about a disease with additional evidence available from clinical and pre-clinical studies and decide which genes to prioritize for experimental validation. The selection process is tedious and time consuming. It also relies on deep specialized knowledge, hence the results can be prone to the individual bias. Our goal was to replace such manual triage with a recommendation solution, which could efficiently integrate diverse types of evidence and identify the most promising candidate genes driving drug resistance.

By moving the problem to a recommendation domain we encounter two major challenges. First is the lack of training data. Here we are dealing with a highly specific molecular phenotype of a poorly understood origin, which prevents us from using information on resistance markers relevant for other, even closely related, diseases as training data. Second, unlike a typical recommendation scenario, in our case both explicit and implicit feedback are lacking. This fact limits our ability to gradually train and improve models. Given these constraints we followed a hybrid unsupervised recommendation approach, which relies on content-based filtering. We formalized re-ranking of CRISPR hits as a multi-objective optimisation problem [26], where diverse and conflicting types of evidence supporting gene’s relevance are mapped to objectives. During the optimization procedure feasible solutions (genes) are identified and compared until no better can be found. A crucial component of such framework is a set of hybrid features. Each feature represents a distinct type of evidence, such as literature support, clinical and pre-clinical evidence.

Along with the purely biological features, our recommendation system relied on data derived from a specially constructed heterogeneous biomedical knowledge graph [27]. Knowledge graphs provide a convenient conceptual representation of relationships (edges) between entities (nodes). In the recommendation context knowledge graphs gain popularity as a way to introduce content-specific information and also to provide explanations for the resulting recommendations [28]. In addition, graph-based recommendations were shown to achieve higher precision and accuracy compared to alternative approaches [29–31]. We used graph structural information together with graph-based representations to express relevance of a gene in the resistance context. Our assumption was that by combining graph-derived features with clinical ones we could discover unobvious genes that drive drug resistance in lung cancer.

In summary, in this study we explored how a question of finding drivers of secondary EGFR TKI resistance could be addressed as a recommendation problem. We demonstrate that a hybrid recommendation system based on multi-objective optimisation approach can be used to re-rank CRISPR hits in the context of secondary drug resistance. The proposed framework, together with an automated feature generation flow and interactive re-ranking interface, helped to reduce gene hit prioritisation time from months to a few minutes. Moreover, it helped to identify resistance candidates that would potentially have been missed by either inherent expert bias or limited integration of experimental outcome with prior knowledge.

## Results

### Re-ranking of CRISPR hits can be approached as multi-objective optimization

We framed re-ranking of CRISPR hits as a multi-objective optimization problem. In this setting, diverse lines of evidence that support gene’s relevance are treated as multiple objectives (Figure 1). In other words, the formal goal is to simultaneously optimize *k* objectives, reflected in *k* objective functions: *f*_1_(*x*), *f*_2_(*x*), …, *f*_*k*_(*x*). Individual functions form a vector function *F* (*x*):

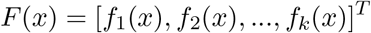

where *x* = [*x*_1_, *x*_2_, …, *x*_*m*_] ∈ Ω; *x* represents the decision variable, Ω - decision variable space. Therefore, multi-objective optimization can be defined as minimization (or maximization) of the objective function set *F*(*x*). With multiple competing objectives a singular best solution often cannot be found. However, one can identify a set of optimal solutions based on the notion of Pareto dominance [32]. A solution *x*_1_ dominates solution *x*_2_ if the following two conditions are true:

**Figure 1:**
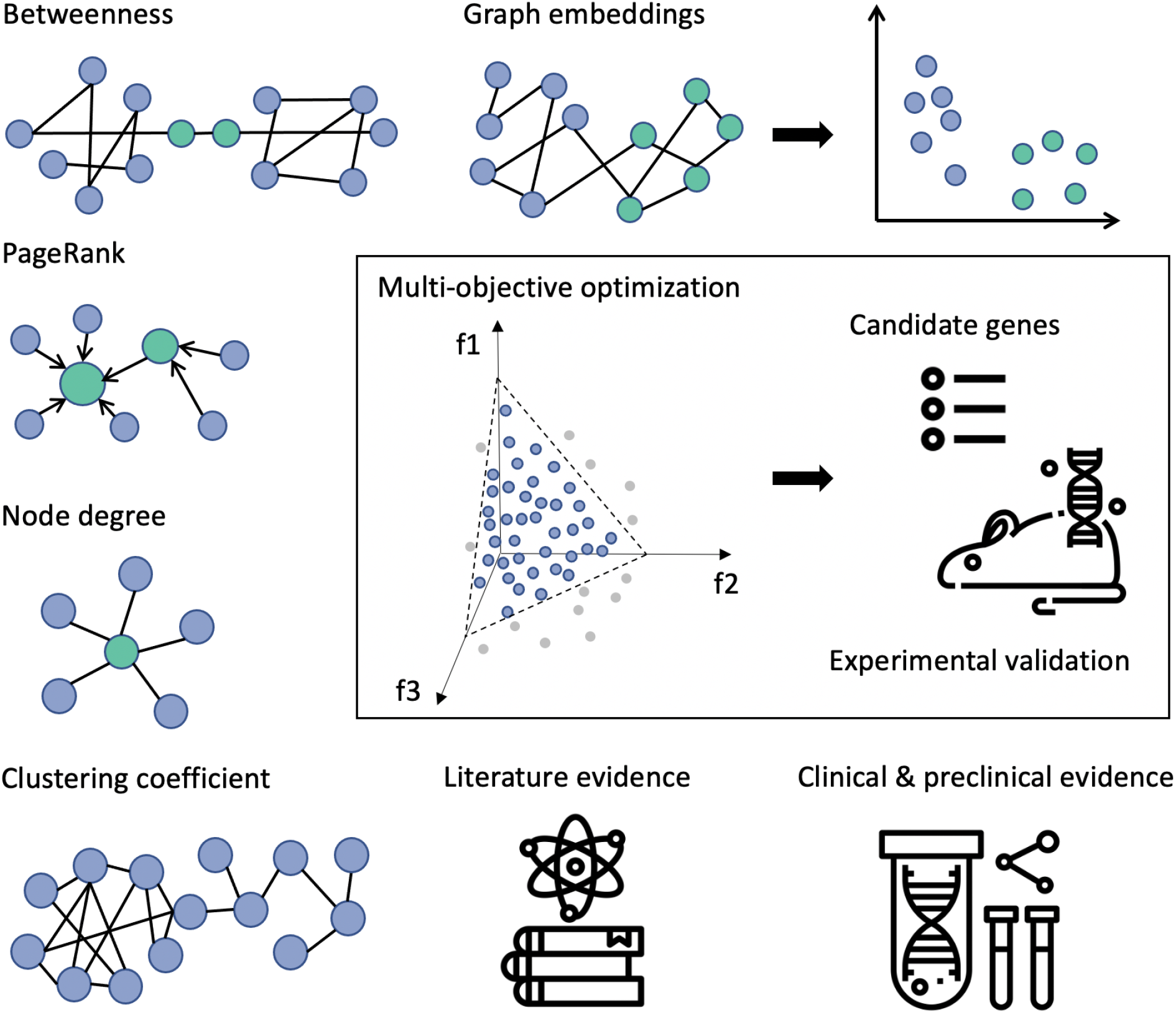
Hybrid recommendation system takes into account diverse types of evidence to suggest promising drivers of drug resistance in NSCLC. The evidence from specially built knowledge graph, literature, clinical and pre-clinical datasets is aggregated and formalized as objectives in multi-objective optimization (MOO) task. Recommended solutions (genes) represent the optimal trade-offs between the conflicting objectives. A subset of recommended genes is passed for the experimental validation.

- solution *x*_1_ is not worse than *x*_2_ according to all objectives;
- solution *x*_1_ is strictly better than solution *x*_2_ according to at least one objective.

If both conditions are true, we can say that *x*_1_ dominates *x*_2_, which is equal to *x*_2_ being dominated by *x*_1_. In other words, dominant solutions can not be improved any further without compromising at least one of the other objectives. A set of such dominant solutions forms a Pareto front, which combines the best trade-off combinations between competing objectives. Therefore, by computing Pareto fronts on diverse sets of objectives defined based on CRISPR screen data and additional supporting evidence we can narrow down the number of promising markers of EGFR TKI resistance (Figure 1).

### A hybrid set of features supports recommendation system

To support the recommendation system we assembled a hybrid set of rich features (Figure 1, Supplementary Table 1), with an idea that each feature represents an objective. The selected features were relevant for EGFR inhibitor resistance in NSCLC and corresponded to distinct lines of evidence. Key feature types and rationale to consider them for re-ranking of CRISPR hits are summarized below.

#### CRISPR

CRISPR screen data served as a starting point for re-ranking. In this study we relied on screens that were set-up to resemble clinical treatment scenarios for *EGFR* mutant lung cancer, using NSCLC cancer cell lines harbouring *EGFR* mutations commonly found in patient populations and where those cell lines were treated with 1^st^ or 3^rd^ generation EGFR inhibitors. In total we identified a starting list of >1000 candidate drug resistance genes [33] that were labelled as significant after the screen analysis. We further aggregated CRISPR data by computing consistency metrics, which reflected stability of a gene’s performance across experimental conditions. Normally genes showing consistent behaviour in multiple relevant conditions, e.g related cell lines or treatments, are ranked higher by domain experts. Altogether, seven consistency-based features were incorporated in the feature set: 1) three features based on loss-of-function part of the screen; 2) three features based on gain-of-function part of the screen; 3) a summary metric reflecting overall consistency in the full screen (Table 1).

### Literature-based metrics

Literature search is routinely used as a first step to confirm experimental findings and to find support for a potential mechanistic hypothesis. For the EGFR inhibitor resistance problem we were primarily interested in the overall literature support for a gene. As a proxy of literature support we calculated the total number of publications that mention a gene in a relevant context, such as *cancer, resistance, EGFR, NSCLC*. Conveniently, the same exact metric when reversed can be interpreted as novelty of a particular target. To extract literature mentions we analysed a total corpus of >180,000 PubMed papers published between 2000 and 2019. We included aggregated literature metrics, based on two terms of interest: EGFR and NSCLC. For each gene we computed the number of papers that mention a gene together with one of these terms (Supplementary Table 1).

### Graph-derived features

In this study we used a custom knowledge graph (KG) as a source of side information for the recommendation system. Our KG contained 11 million nodes and 84 million edges and was composed of 37 public and internal datasets, such as Hetionet, OpenTargets, ChEMBL and Ensembl [27]. In general, patterns of interactions between biological entities captured by knowledge graphs can be translated into features and consumed by recommendation systems in a number of ways (Figure 1, Supplementary Table 1). One way is to compute features directly on the graph. This includes metrics such as node degree — reflecting the importance of a node; PageRank — a measure of node’s popularity [34]; betweenness — a way to detect the amount of influence a node has over the flow of information in the graph. An alternative approach involves projecting the graph into a low-dimensional space, so that every node is transformed into its vector representation — embedding. Embeddings capture critical structural properties of the graph [35], so that the nodes that were close in the graph also remain close in the embedding space. In this study we computed distances in the embedding space between each gene and two key entities of interest: ‘EGFR’ and ‘NSCLC’. The assumption is that genes most relevant to the EGFR TKI resistance phenotype should be close to either lung cancer or EGFR gene nodes.

### Clinical enrichment scores

To ensure the recommendation system captures clinical evidence, we included genomic data from osimertinib-treated EGFR-mutant lung cancer patients in the feature set. We prioritised four clinical trials: AURAext[36], AURA2[37], AURA3[21], and FLAURA[22]. The prevalence of genomic alterations in non-responders vs responders across 355 patients treated with osimertinib were calculated and included as ’clinical enrichment score features’ to the feature set (Supplementary Table 1).

### Tractability and gene essentiality

Traditionally drug resistance in cancer is addressed by developing compounds or combination therapy that modulates activity of its key driver genes (targets). When a target is prioritised for drug development one needs to ensure that: 1) a gene is tractable in principle, i.e it is shown or predicted to bind to commonly used drug modalities with high affinity; 2) a gene should not be essential, since knockout of an essential gene can be detrimental to other cells in the organism, not just the tumour ones. To support the first consideration, we included bucket tractability estimates [38] for three modalities: antibodies, small molecules and other modalities (enzyme, oligonucleotides, etc). In support of the second consideration we integrated DepMap [39] essentiality estimates.

In summary, the final hybrid set contained 25 rich features, supporting diverse criteria taken into consideration during validation of CRISPR hits by domain experts (Supplementary Table 1). The hybrid set was also augmented by graph-derived features and literature-based metrics.

### Interactive interface allows experts to re-rank CRISPR hits

So far, we have defined a basic model for multi-objective optimization and demonstrated how to build a hybrid set of features to support re-ranking of CRISPR hits in the EGFR TKI context.

In the real-world scenario, decision making can be both iterative and subjective. A choice of a particular set of objectives and the direction of optimization for the same variable varies from expert to expert. Each combination of objectives and corresponding directions for optimization might result in a different shape of Pareto front, therefore — in a different set of top recommended genes.

To accommodate diversity of opinions and to enable domain scientists to explore complex tradeoffs between the objectives we built an interactive application — skywalkR https://github.com/AstraZeneca/skywalkR (Figure 2). SkywalkR is a Shiny app [40], which operates on top of the pre-assembled hybrid feature set (see Supplementary Table 1). SkywalkR app combines diverse facets of knowledge to guide re-prioritization of CRISPR hits for experimental validation. In addition, it allows domain experts to explore various trade offs between objectives. Thereby, it stimulates exploration of possibilities, highlights gaps in the existing knowledge and motivates to adjust expectations about optimal solutions.

**Figure 2:**
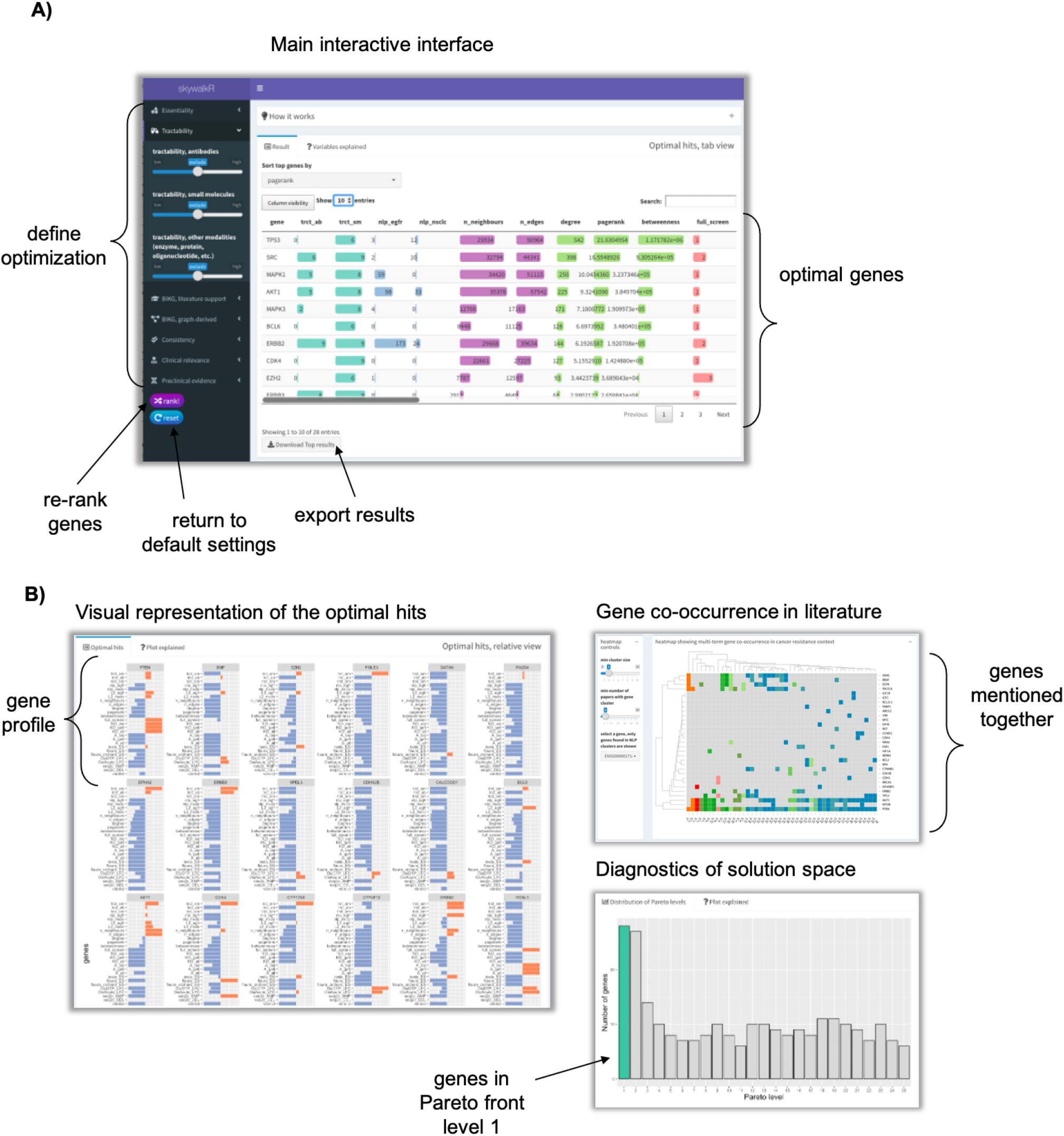
SkywalkR interactive interface allows users to re-rank CRISPR hits based on various combinations of objectives. **A**) On the side bar panel each objective is represented with a slider. Users can decide which objectives to include in the optimization and can also specify direction of optimization (minimize or maximize). **B**) Additional tools to explore the results. Relative view shows profiles of recommended genes. Bar plots demonstrate standardized values across objectives and top recommended genes. Co-occurence heatmap demonstrates clusters of genes frequently mentioned together in EGFR TKI resistance context.

Automated engineering of rich features coupled with multi-objective optimization realized through skywalkR interactive interface dramatically reduced the time required for gene prioritization from a few weeks to minutes.

### Evaluation demonstrates majority of top recommendations labelled as credible by the experts

To evaluate the recommendation framework against the expert opinions we fixed a default set of preferences. Preferences were defined by a combination of selected objectives and corresponding directions of optimization. The set of defaults was chosen to mimic the process of CRISPR hit validation by domain experts. The resulting list contained 36 recommended genes (Figure 3). To collect opinions on the list from the domain experts, we set up an interactive evaluation task with Prodigy [41]. Five independent experts assigned each of the recommended genes to one or more predefined categories: 1) known, resistance marker; 2) novel, but credible hit; 3) novel, not credible hit; 4) not novel, not credible hit. Here ’credible’ referred to tractability and meant an existence/absence of a clear path to biological validation.

**Figure 3:**
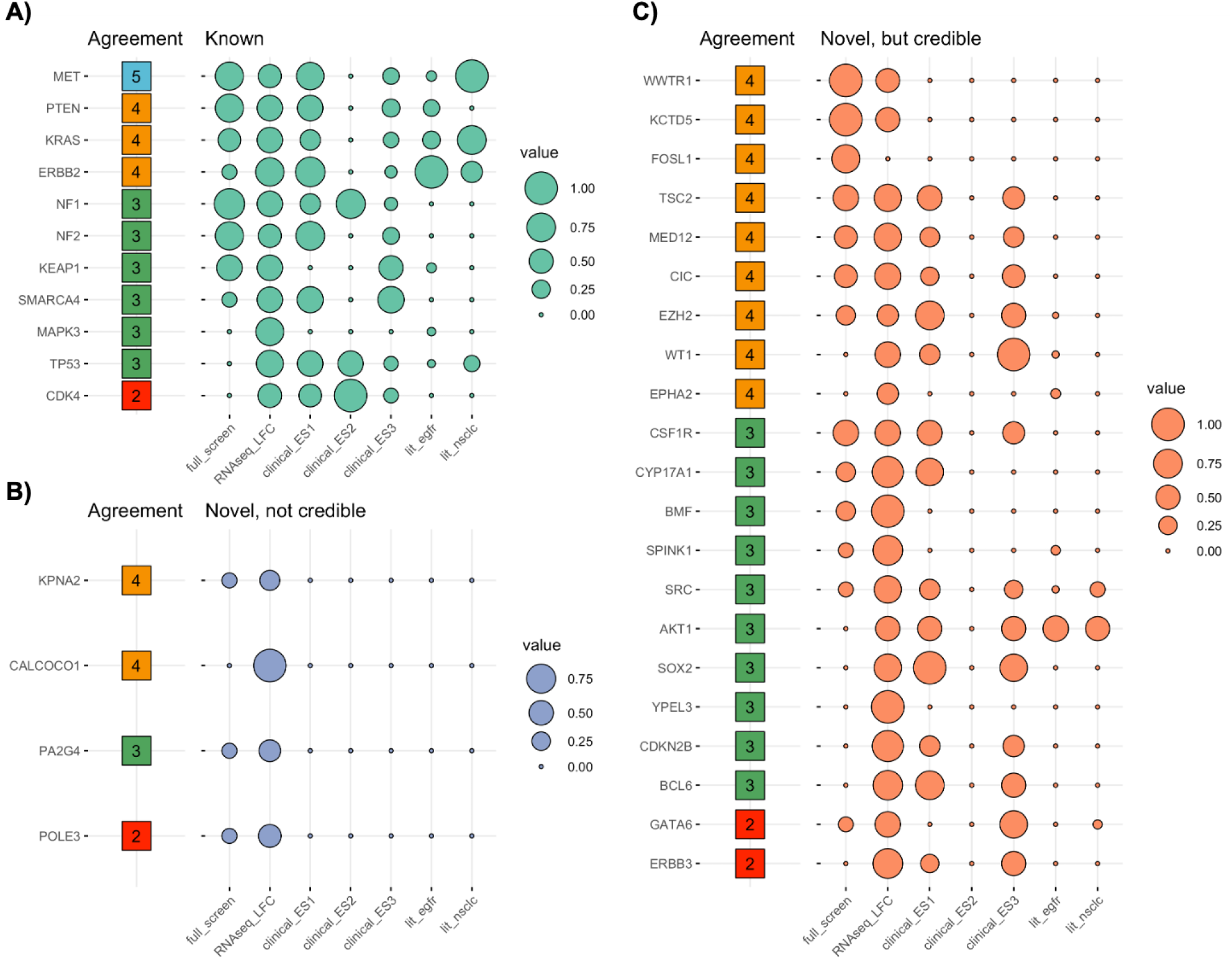
Evaluation of top recommended genes by five independent experts indicates that majority of genes can be classified either as ‘known’ **A)** or ‘novel, but credible’ **B)** categories. Four genes were labelled as ‘novel, not credible by the experts’ **C)**. Here credibility refers to the absence of clear path to validate predictions experimentally. Features used to generate predictions: *full_screen* – overall consistency across all conditions in the CRISPR screen; *RNASeq_LFC* – log2 fold change from internal RNASeq study; *clinical_ES1, clinical_ES2, clinical_ES3* – enrichment scores from clinical studies where resistant patients were compared against responders; *lit_EGFR, lit_NSCLC* – co-occurrence estimates from the literature. *Agreement* column indicates the number of experts assigning a certain label to a gene. Size of a bubble reflects value normalized across the full set of features for all genes.

Despite the expected discrepancies between the expert opinions, the majority of the recommended genes (89%) were classified as either ‘novel, but credible hit’ or ‘known resistance marker’ (Figure 3). To consolidate opinions we assigned the most frequent label to each gene (best label). This resulted in three major categories: ‘known’, ‘novel, but credible’ and ‘novel, not credible’. To determine underlying data structure that supported separation between the three labels, we analysed values corresponding to seven objectives included in the default preference. For easier comparison values were standardised (Figure 3). Genes labelled as ’novel, not credible’ were clearly separated from the remaining genes on the basis of low values of all objectives across the board, except the log fold change values from the RNA-Seq study 3. This analysis suggests that in general the experts tend to prioritize genes supported by several lines of evidence.

### Shapley values indicate the high impact of CRISPR-derived features

To further estimate what was the impact of each of the objectives on the expert decisions, we calculated Shapley values [42], a game-theoretic technique used to explain output of machine learning models. For this analysis we reduced the problem to a binary classification task, where a gene is either selected by an expert or not. To assign positive labels we used a set of 100 genes prioritized to a secondary CRISPR screen [33] and trained two random forest models: 1) based on a default subset of features; 2) based on the full set of rich features, including clinical, pre-clinical, literature, CRISPR and graph-derived categories (Figure 4).

**Figure 4:**
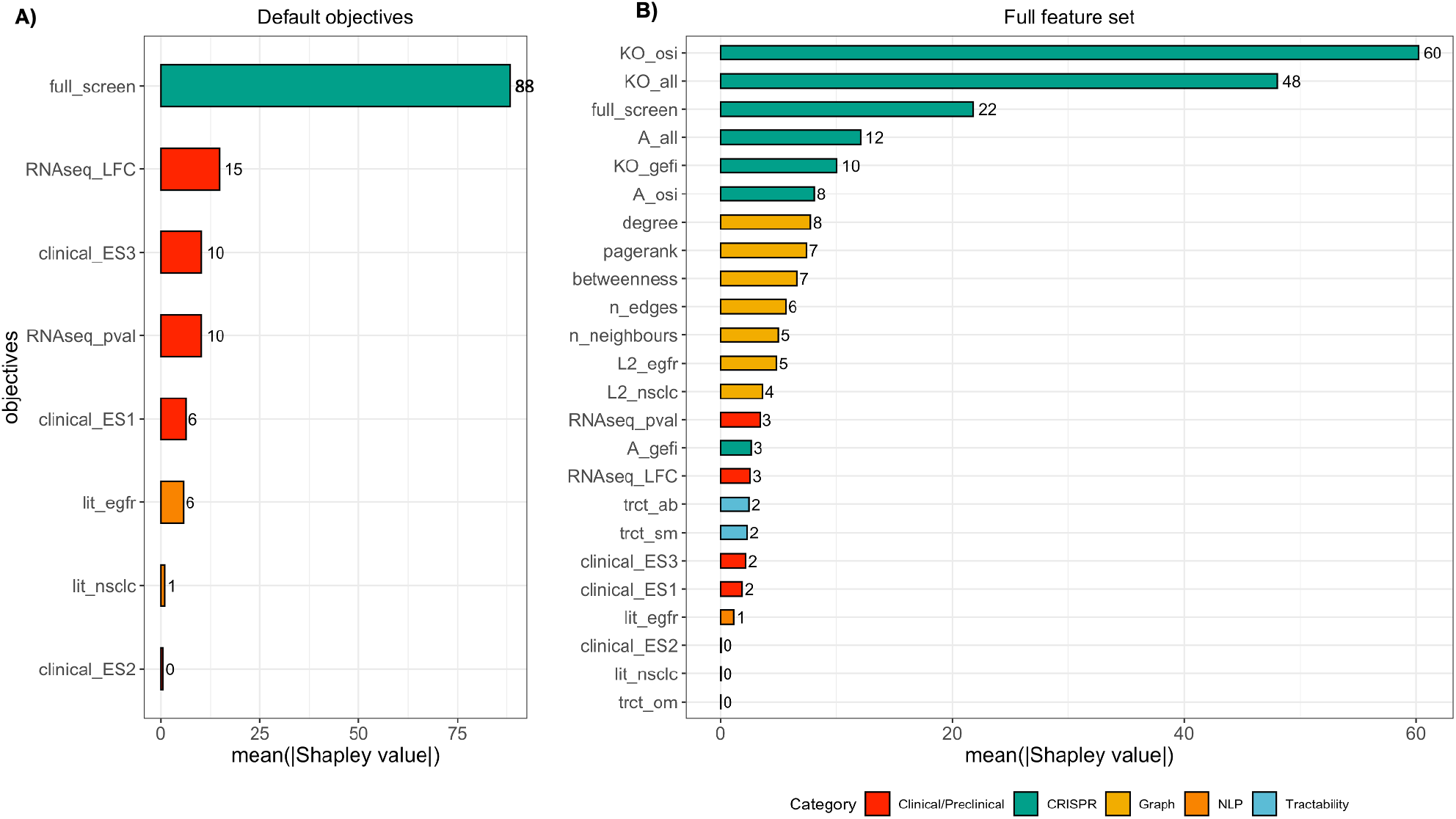
Shapley values reflect relative importance of features that differentiate relevant hits from non-relevant. We modelled the task as a binary classification problem, where a gene can be labelled as either relevant or not. A list of genes prioritised by domain experts for secondary screens was used as a set of labels. The analysis was run using a subsection of default objectives **A)** and repeated on the full set of objectives **B)**. In both cases features associated with consistency in CRISPR screens appear to have the greatest impact on classification.

The resulting Shapley values indicate that in general, CRISPR-derived features have the largest impact on gene classification. When the default set of features was tested, the composite CRISPR screen consistency variable accounted for the most impact on classification 4. Interestingly, when a full set of features was used to train the model, CRISPR-derived features remained decisive, followed by graph-derived features. This suggests that graph-derived features though not routinely used in manual triage, could provide a valuable insight in the overall relevance of a gene, especially when combined with context-defining experimental data, such as CRISPR.

### Pathway enrichment analysis captures known and novel EGFR resistance mechanisms among top recommended genes

To link the prioritised hits to known EGFR biology, we performed pathway enrichment analysis. It captured pathways related to resistance such as ’mechanisms of resistance to EGFR inhibitors in lung cancer’ and ’LKB1 signaling pathway’ (Table 2). Occasionally enrichment results are redundant and top terms may be similar to each other thus carrying little new information. Therefore we also performed crosstalk analysis (Supplementary Figure 6) which confirmed ’mechanisms of resistance to EGFR inhibitors in lung cancer’ as the top enriched pathway.

### Experimental validation demonstrates that deregulation of a subset of recommended genes mediates resistance phenotype

To further validate a subset of recommended genes we experimentally investigated their direct impact on osimertinib resistance. We manipulated expression of four recommended genes (MET, PTEN, NF1 and KCTD5) in *EGFR* mutant NSCLC cell lines PC-9 and HCC827, both sensitive to osimertinib (Figure 7, A).

Our expectation was that deregulation of PTEN, NF1 and KCTD5 should mediate a stable resistance phenotype, since these genes were shown to negatively regulate MAPK or PI3K/AKT signalling [43–45] – known drivers of EGFR TKI resistance. To test this hypothesis we established a flow cytometry based long-term competition assay (Figure 5, B). The assay showed that after 14 days of co-culture perturbation of NF1, PTEN or KCTD5 expression did not affect proliferation compared to control cells. However, when treated with osimertinib NF1 or PTEN KO caused a fitness advantage, measured as a two to three fold increase in proliferation (in PC-9 or HCC827, respectively) compared to control cells (Figure 5, C). These results indicate stable and profound osimertinib resistance mediated by PTEN and NF1.

**Figure 5:**
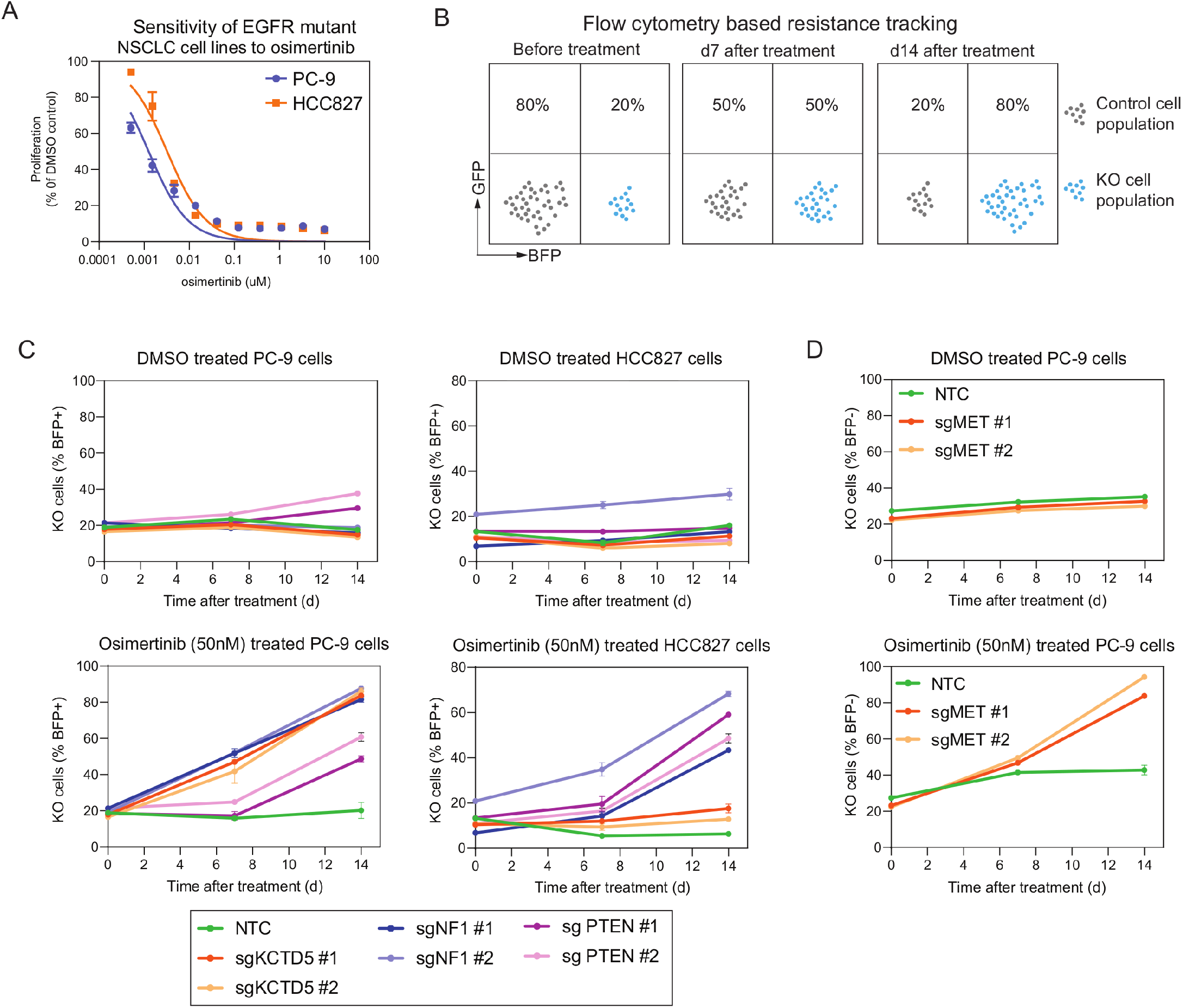
Validation of resistance genes proposed by recommendation system. **A)** Proliferation assays confirm osimertinib sensitivity of EGFR-mutant NSCLC cell lines PC-9 and HCC827 after 5 days of treatment. **B)** Resistance to osimertinib by target gene KO was measured in flow cytometry based competition assays.Increase in percentage of KO cells compared to control cells was considered as resistance effect to osimertinib. **C)** Confirmation of resistance to osimertinib in *EGFR* mutant NSCLC KO cell line models. Effect of KO of three genes (KCTD5, NF1 and PTEN) on osimertinib resistance was tested in PC-9 and HCC827 cell lines. Proliferation of cells in DMSO or osimertinib (50nM) was tracked over 14 days and percentage of BFP+ KO cells was plotted. D) Activation of MET drives resistance to osimertinib in *EGFR* mutant PC-9 cells PC-9 cells were engineered to express the dCas9-VP64 SAM CRISPR activation system [72].

Recent studies indicate that amplifications of MET [23] are often associated with an overexpression of this receptor tyrosine kinase resulting in bypassing of EGFR signals [46]. To validate relevance of MET to osimertinib resistance we activated its expression in PC-9 cells (Figure 7, B) and compared proliferation in control (DMSO) and drug treatment. As expected, overexpression of MET did not alter cell proliferation under control conditions but was substantially elevated when cells were treated with osimertinib compared to control modified cells (Figure 5, D).

Together these findings indicate that three out of four selected recommended genes - PTEN, NF1 and MET have initial experimental evidence indicating their involvement in osimertinib resistance.

### Recommendation approach identifies novel candidate genes involved in EGFR TKI resistance

In addition to known EGFR TKI resistance markers, and validated markers discussed above, our method also identified several novel markers of osimertinib resistance (Figure 3). Here we define ’novel’ as genes having no publications that directly associate them to EGFR in the context of resistance.

Two of these novel hits, FOSL1 and BCL6, have been since shown to be involved in key molecular bypass mechanisms for EGFR-TKI resistance [47], [48]. FOSL1 has been shown to play a key role in the cross-talk between MEK and Hippo pathways. Misregulation of MEK pathway in driving tumour growth is well established [49], and key Hippo pathway members (YAP, TAZ) have been implicated in NSCLC [50]. Pham et al.[47] showed that a significant decrease in FOSL1 expression was observed when YAP1-amplified Detroit562 cells were treated with a combination of YAP1 knockdown and Cobimetinib (MEKi), but not for either treatment. BCL6 plays a key role in mediating core cell functions such as antiapoptosis and DNA Damage recognition and has been shown to play a key role in NSCLC [51]. Tran et al. [48] showed that inhibition of BCL6 in NSCLC cell lines conferred sensitivity to Gefitinib. Further experiments also showed that targeting BCL6 and EGFR as a combination showed significant synergy. Altogether these observations indicate that our recommendation approach suggested not only well known resistance markers, but was also able to identify novel and promising drivers of resistance.

## Discussion

In this study we explored how an open-ended biological question — discovering drivers of drug resistance in lung cancer — can be approached as a recommendation problem. The current protocol to find resistance markers starts with high-throughput CRISPR screens, followed by a lengthy and time-consuming manual triage which requires specialised knowledge. Our goal was to replace the manual triage with a recommendation solution that outputs genes potentially driving the resistance. Undoubtedly, such a biomedical setting is not a traditional area of application for recommendation systems. Still, the need to find a small number of relevant genes amongst prohibitively large set of possibilities constitutes a typical recommendation task. How to approach it?

Common recommendation problems are often solved with collaborative filtering [52], content-based [53] or hybrid solutions [54]. The idea behind collaborative filtering is to predict preferences of a user for an item based on weighted preferences of other users [52]. This approach was demonstrated to work successfully to recommend conceptually complex entities, such as movies, without any additional information about the entity itself [55]. However, to work accurately collaborative filtering requires either large datasets of users actively rating items or a substantial amount of historical data. Due to the lack of user interactions data, or its equivalent, collaborative filtering is not directly applicable to the CRISPR hit re-ranking task.

On the other hand, content-based approaches recommend items exclusively based on the properties of an item and do not require user interaction data. Therefore, this class of methods seems more suitable for our case. Such content-based recommender would require a set of features, describing properties of a gene relevant for secondary resistance in lung cancer. The disadvantage of content-based approaches is their reliance on similarities between items to make a recommendation. They require either a point of reference or more broadly — training data to make predictions. In the CRISPR hit re-ranking case training data is lacking mainly because a clear conceptual representation of a ***good resistance hit*** is also lacking. The closest analogue of training data in this case would be a weakly labeled dataset [56] created based on information about genes implicated in drug resistance in related types of cancer under similar treatments. It still remains to be proved, however, if a notion of resistance is directly transferable between very specific cancer contexts. Moreover, in this study we focus on discovery of novel markers of drug resistance, that have never been explored before. Hence it won’t make sense to base recommendations on a handful of genes that are already known to be implicated in drug resistance.

In short, CRISPR hit re-ranking task does not immediately fit into classic recommendation frameworks. Complexity of the biological context, lack of insight about mechanisms of resistance, coupled with absence of training data and user feedback make it difficult to build a fully fledged recommendation solution capable of reasoning about drug resistance.

Devoid of user feedback and training data we approached CRISPR re-ranking problem as a decisionmaking task. We formalized the problem as a multi-objective optimization with the goal to find solutions defined by optimal trade-offs between novelty, druggability, clinical and pre-clinical relevance. This approach operates in an unsupervised domain, where a perfect solution is unknown. Still we can leverage domain expertise to loosely define a profile of a ’good resistance hit’. This can be formalized by including/excluding objectives, setting directions of optimization and setting constraints on the objectives themselves.

Similar to the manual triage, re-ranking of CRISPR hits through multi-objective optimization can be performed iteratively. By experimenting with diverse sets of objectives one could explore plausible solutions given the constraints of underlying data. We recognised such iterative decision making could benefit from an interactive interface. Therefore we developed a Shiny app, SkywalkR, that accounts for diversity of opinions and allows the users to explore the resistance space more efficiently. The app allows the experts to compose custom optimization preferences and re-rank the list of hits based on them. The overall technical solution includes: 1) an automated pipeline that generates features; 2) an interactive interface that runs on top of multi-objective optimization model and has access to the features. This setup helped to dramatically reduce the time required to prioritise CRISPR screen hits from the usual weeks-months to a few minutes.

The key component of our recommendation system is a hybrid set of features, which is tailored to the EGFR inhibitor resistance context. Along with the usual types of evidence used by the experts during the manual triage, such as clinical and pre-clinical evidence, we included features derived from a custom heterogeneous biomedical knowledge graph. The rationale behind it is that structural properties of the knowledge graph can express relevance of a gene, even if a direct proof of its association with resistance does not yet exist in the literature or clinical/pre-clinical sources. In other words graph-derived features could aid discovery of novel or unexpected drivers of drug resistance.

To determine if our recommendation approach produces meaningful results we developed a hybrid strategy. Recommendations were initially evaluated by domain experts, followed by targeted experimental validation of a few promising genes. The *in silico* evaluation demonstrated that majority (89%) of suggested genes were classified by independent experts as credible and/or novel. The remaining genes (12%) were classified by the experts as ’novel but not tractable’, meaning there was not a clear path for biological validation yet. To complement the evaluation we picked four genes (MET, PTEN, NF1 and KCTD5) for experimental validation. We demonstrated that deregulation of these genes indeed mediated stable resistance phenotype. Though the exact mechanism of this is effect remains to be investigated, our experiments proved that recommended genes are implicated in secondary resistance. In summary, hybrid evaluation demonstrates that our recommendation approach not only produces relevant results, but also does it in a fraction of a time compared with the manual triage.

Though the initial results are promising we recognise that overall our recommendation approach has a number of limitations and areas of improvement. First of all, when applying the multi-objective optimization approach to the CRISPR hit triage problem, there is a risk to consider an excessive number of objectives/lines of evidence. The more objectives we take into account, the broader and more topologically complex Pareto fronts can become. This effect limits our ability to unambiguously select a small set of optimal solutions. A few strategies can help to overcome this problem: 1) select a small number of the most important objectives relying on the domain knowledge; 2) multiple objectives can be combined into a single one using scalarization techniques [57]; 3) introduce adaptive weights to individual objectives based on the domain knowledge and the notion of the relative importance of each type of evidence [31]; 4) multi-objective optimization can be performed in consecutive stages on a sub-selection of objectives, similar to Markov decision process [58]. Some of the above approaches could be combined, for example scalarization and adaptive weights. Worth mentioning that all of the listed strategies rely on domain knowledge. Perhaps, they can be viewed as a progressive way to translate individual researcher bias into a formal model.

Another limitation we faced in this study is difficulties with assessing the accuracy of our recommendations. Primarily it stems from the lack of clear notion of ’good resistance hit’. In addition we could not rely on user feedback to gradually evaluate and improve predictions. The ultimate source of truth in our case is experimental validation, where a role of a gene in driving resistance phenotype could be tested *in vitro* or in animal models. At the moment large-scale validation experiments are not feasible since they are costly and take long time to perform. However in an ideal scenario, experimental output can become an equivalent of user feedback and be used to improve predictions of biomedical recommendation systems.

In summary, accumulation of large amounts of biomedical data coupled with the need to comprehend and reason about it makes biomedical applications an attractive field for recommendation techniques. However direct translation of traditional recommendation approaches to the biomedical domain is not always trivial. Specifics of the problem space and complexity of biological systems call for efficient recommendation solutions that could operate in unsupervised or weakly supervised settings. We believe that wider adoption and systematic use of recommendation system to solve biological problems bear a potential to transform biomedical research and drug discovery.

## Methods

### Pooled drug CRISPR screen data

The primary data for pre-clinical drug resistance to EGFRi was generated from pooled, genome-wide CRISPR loss-of-function (LoF) and gain-of-function (GoF) studies, analogous to [59]. In the screens every gene in the genome was knocked out or over expressed and the impact on osimertinib resistance was assessed based on a treatment-control comparison of guide RNA prevalence. The screens recapitulate clinical treatment scenarios for *EGFR* mutant lung cancer in that the cell line models contain activating mutations of EGFR. As described in [33] the cell lines were treated with EGFR inhibitors for 3 weeks to allow hit identification of genes driving resistance when knocked out or overexpressed. In total we identified a starting list of >3000 resistance genes as hits to be re-ranked. Hits were defined according to [60].

### Graph-derived features

The biomedical knowledge graph was built as described in [27]. Embeddings were calculated based on the full graph using RESCAL algorithm [61]. L2 distance from human gene nodes to ’EGFR’ and ’NSCLC’ nodes was calculated using Faiss package [62]. Full graph was also used to calculate descriptive network metrics such as node degree and number of unique neighbours connected to a node.

To make graph-derived metrics more relevant to mechanistic explanation of EGFRi resistance, we further focused on a protein-protein interaction (PPI) subgraph. PPI subgraph was defined based on the ‘interacts’ edges from HetioNet [63, 64]. PPI subgraph was used to calculate PageRank [34] and betweenness [65] metrics for each gene node.

### Literature-based features

To estimate overall literature support for a gene’s involvement in EGFR inhibitor resistance we analysed a corpus of PubMed and PMC articles as well as Springer bio-classified data published between 2000 and 2019. The corpus was further restricted to a set of 185299 publications relevant to cancer and/or secondary drug resistance. The search was performed on the title, abstract and full text. Next, two key terms were identified - ’EGFR’ and ’NSCLC’. For each of the target terms we computed the total number of papers that mention a given human gene and one of the key terms of interest together. The resulting two summary metrics were included in a hybrid feature set and exposed through the skywalkR interface.

In addition to single-gene metrics we computed multi-term gene co-occurrence in previously defined (cancer and drug resistance) context. The idea behind this analysis was to discover sets of genes that tend to co-occur together in EGFR inhibitor resistance context. Frequent co-occurrence of gene combinations across publications could indicate a strong link between genes within a set and suggest a potential mechanism driving secondary resistance. Gene co-occurrence matrix was used to build an interactive heatmap in the skywalkR app.

### Clinical enrichment features

Osimertinib is an irreversible EGFR inhibitor that selectively targets the EGFR T790M mutation [66]. We aggregated data from Osimertinib-treated patients across four clinical trials - AURAext, AURA2, AURA3, and FLAURA. AURAext is a phase II extention of the AURA trial with an 80mg/day Osimertinib dose administered to Non-small Cell Lung Cancer (NSCLC) patients [36]. AURA2 is a Phase 2 single arm clinical trial for NSCLC patients with advanced disease who progressed on previous treatment with EGFR Tyrosine Kinase Inhibitor (TKI), and also carry the EGFR T790M mutation [37]. AURA3 is a Phase 3 randomised study comparing the efficacy of Osimertinib vs platinum-based chemotherapy in advanced NSCLC patients who have progressed on prior treatment with EGFR TKI. These patients also carry the EGFR T790M mutation [21]. FLAURA is a Phase 3 clinical trial for first-line Osimertinib treated advanced NSCLC patients vs other EGFR TKI standard-of-care (SoC) treatments [22].

335 patients treated with Osimertinib across the 4 trials were analysed. Sequencing data from Guardant Health and FMI gene panels were utilised to identify genetic alterations. Patients who were classified with the RECIST criteria of Partial Responder/Complete Responder AND PFS > 6mos were classified as responders, and enrichment of genetic alterations in responders vs non-responders was calculated. As individual trials utilised different clinical gene panels, the enrichment metrics were kept trial-specific and no aggregation across trials was performed. This ’enrichment score’ was used as a feature for multi-objective optimisation.

### Tractability

To estimate druggability of candidate genes we relied on tractability scores from OpenTargets https://github.com/melschneider/tractability_pipeline_v2, [38]. Tractability scores for three modalities were included: antibodies, small molecules and other modalities (e.g. enzyme, protein, oligonucleotide, etc.). For convenience buckets were reversed, so that the highest druggable bucket corresponded to the highest numeric score 10. Reversed tractability metrics were exposed through the skywalkR app.

### Gene essentiality

Information about whether the gene is common essential or not was retrieved from DMC DepMap [67]. We flagged a gene as “essential” if inhibition of target gene results in reduced viability for 90% of cell lines used in DepMap. Otherwise a gene was flagged as “nonessential”.

### Models, implementations

Pareto fronts were computed based on the implementation from the rPref R package [68]. To build binary classifiers for gene labels we used fast implementation of random forest from ranger package [69]. Shapley values were calculated using fastshap package [70].

### Pathway enrichment analysis

We used MetaCore to perform enrichment analysis. Genes were matched to possible targets in functional ontologies of MetaCore. The probability of a random intersection between a set of IDs the size of target list with ontology entities is estimated in p-value of hypergeometric intersection. The lower p-value means higher relevance of the entity to the dataset. Canonical pathway maps represent a set of signaling and metabolic maps for human. All maps are created by manual curators/scientists from Clarivate Analytics relying on published peer-reviewed literature. To adjust enrichment results we additionally performed crosstalk analysis.

### Generation of KO or activation cell lines, plasmids and antibodies

PC-9 and HCC827 cell lines were cultured at 37C and 5% CO2 in RPMI 1640 GlutaMAX media (Gibco, US) supplemented with 10% fetal bovine serum. For generation of KO cell line pools PC-9 and HCC827 cells were transduced with pKLV2-EF1a-Cas9Bsd-W (Addgene ID:68343)[71] to stably express Cas9. Transduced cells were subjected to Blasticidin (Gibco, US) selection. To knockout target genes guide RNAs targeting KCTD5, NF1, PTEN or non-targeting-control (see Table 3) were cloned into pLKV2-U6gRNA5(BbsI)-PGKpuro2ABFP-W (Addgene ID:67974) and Cas9 expressing cell lines were transduced and selected with puromycin (Gibco, US). To activate MET expression in PC-9 cells the three vector based CRISPR SAM system [72] was used. In brief, viroid’s containing open reading frames of dCas9-VP64 and MS2-P65-HSF1 as well as target specific guideRNA expression constructs were purchased (SAMVP64BSTV, SAMMS2HYGV and LV06 respectively, Sigma Aldrich) and used to stepwise transduce and select PC-9 cells. After each transduction cells were subjected to respective antibiotic selection (blasticidin, hygromycin, puromycin). 14 days after transduction whole cell lysates of selected cell pools were analysed by western blot to confirm CRISPR KO or CRISPR activation of gene expression using anti-KCTD5 (proteintech, 15553-1-AP), anti-MET (CST, #8198), anti-NF1(abcam, ab17963), anti-PTEN (CST, #9559), as well as loading control ab.

### Viability assay

Two thousand cells were seeded 24h prior to treatmemt into 96 well plates. Cells were treated with indicated concentrations of DMSO or osimertinib and cultured for 5 days as technical triplicates. Viability was determined by adding CellTiter-Glo Luminescent Cell Viability Assay (Promega, US) according to manufacturer’s protocol and measuring luminescence using a SpectraMax plate reader (Molecular Devices, US). Results are visualised as percent viability of DMSO treated control. Shown are representative results of three biological replicates.

### Flow cytometry based resistance tracking

For studying the effects of gene KO on osimertinib resistance, PC-9 and HCC827 KO cell lines were cocultured with respective Cas9-expressing control cells and treated with DMSO or osimertinib (50nM) for 14 days. Media and drug was replenished every 3-4 days. Fractions of KO cells relative to control cells were determined by measuring BFP co-expression in KO cells via flow cytometry at indicated timepoints. In case of PC-9 CRISPR activation lines, PC-9 control cells were stably labelled with BFP and fractions of unstained CRISPR activation cells relative to BFP-positive control cells were determined by flow cytometry. All experiments were performed as three technical replicates of two independent guideRNAs per gene as biological replicates. FlowJo 10 was used for analysis of FACS data.

## Code availability

skywalkR source code and documentation can be found at https://github.com/AstraZeneca/skywalkR.

## Acknowledgements

We thank David Geleta, Andriy Nikolov, Gavin Edwards, Benedek Rozemberczki and Daniel Barrel for technical and scientific support. We thank Paul Smith, Steven Criscione, Matthew Martin and Aisha Swaih for expert validation of candidate hits. We thank the Early Computational Oncology and Functional Genomics Centre teams for scientific input and feedback.

## Author contributions statement

K.B., U.M., B.S, J.D. and E.P. conceived the study. A.G. designed and implemented the computational framework and analyzed the data. D.P, V.P, M.A and K.B. generated and analyzed the data. M.P. carried out wet lab experiments and analyzed the data. A.G. drafted the manuscript in consultation with K.B, D.P, E.P, M.P. and M.A. All authors reviewed the manuscript.

## 1 Supplementary Figures and Tables

**Table 1: SUPPLEMENTARY.**
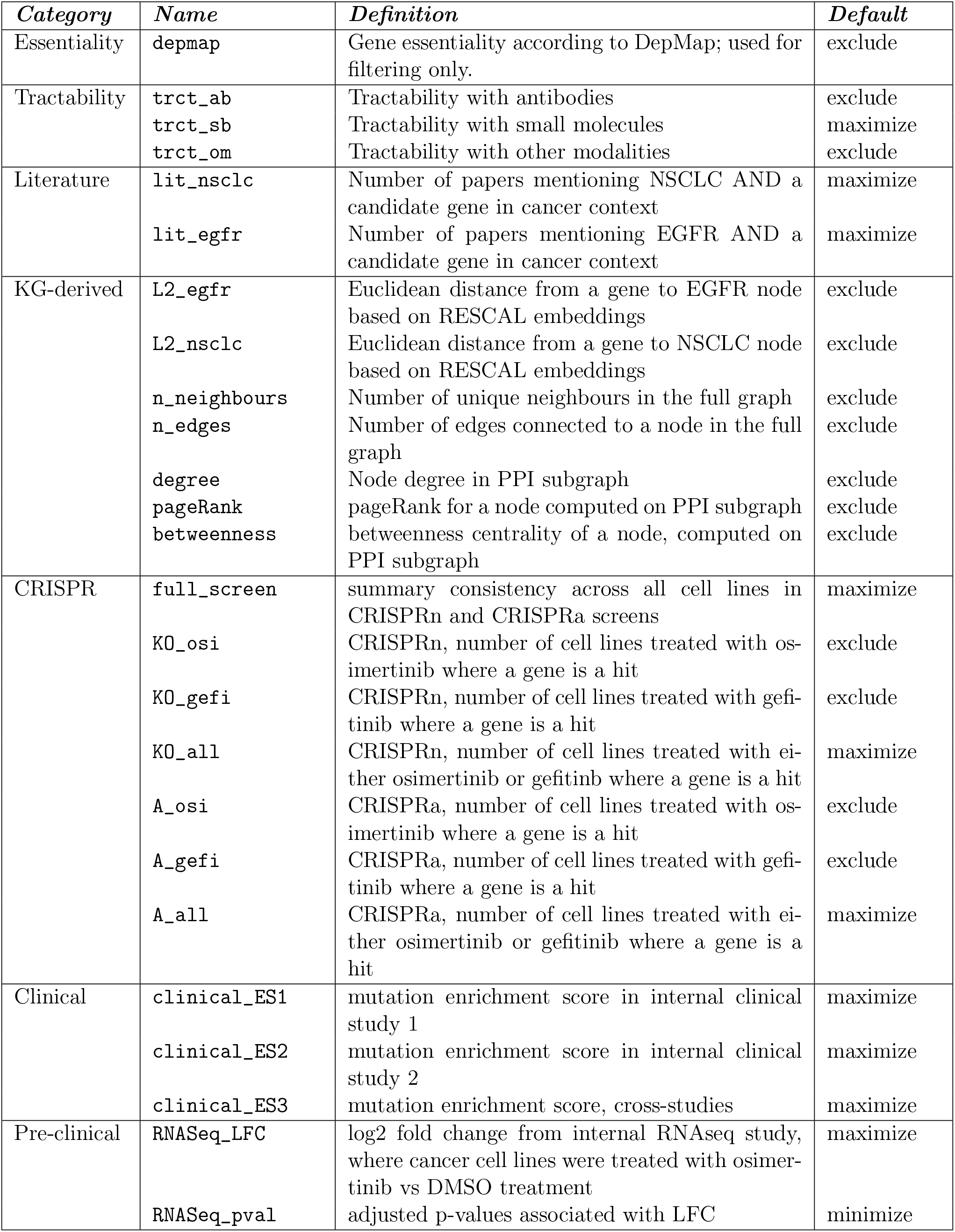
Definitions of features used for multi-objective optimization

| *Category* | *Name* | *Definition* | *Default* |
| --- | --- | --- | --- |
| Essentiality | depmap | Gene essentiality according to DepMap; used for filtering only. | exclude |
| Tractability | trct_ab<br>trct_sb<br>trct_om | Tractability with antibodies<br>Tractability with small molecules<br>Tractability with other modalities | exclude<br>maximize<br>exclude |
| Literature | lit_nsclc<br>lit_egfr | Number of papers mentioning NSCLC AND a candidate gene in cancer context<br>Number of papers mentioning EGFR AND a candidate gene in cancer context | maximize<br>maximize |
| KG-derived | L2_egfr<br>L2_nsclc<br>n_neighbours<br>n_edges<br>degree<br>pageRank<br>betweenness | Euclidean distance from a gene to EGFR node based on RESCAL embeddings<br>Euclidean distance from a gene to NSCLC node based on RESCAL embeddings<br>Number of unique neighbours in the full graph<br>Number of edges connected to a node in the full graph<br>Node degree in PPI subgraph<br>pageRank for a node computed on PPI subgraph<br>betweenness centrality of a node, computed on PPI subgraph | exclude<br>exclude<br>exclude<br>exclude<br>exclude<br>exclude<br>exclude |
| CRISPR | full_screen<br>KO_osi<br>KO_gefi<br>KO_all<br>A_osi<br>A_gefi<br>A_all | summary consistency across all cell lines in CRISPRn and CRISPRa screens<br>CRISPRn, number of cell lines treated with osimertinib where a gene is a hit<br>CRISPRn, number of cell lines treated with gefitinib where a gene is a hit<br>CRISPRn, number of cell lines treated with either osimertinib or gefitinib where a gene is a hit<br>CRISPRa, number of cell lines treated with osimertinib where a gene is a hit<br>CRISPRa, number of cell lines treated with gefitinib where a gene is a hit<br>CRISPRa, number of cell lines treated with either osimertinib or gefitinib where a gene is a hit | maximize<br>exclude<br>exclude<br>maximize<br>exclude<br>exclude<br>maximize |
| Clinical | clinical_ES1<br>clinical_ES2<br>clinical_ES3 | mutation enrichment score in internal clinical study 1<br>mutation enrichment score in internal clinical study 2<br>mutation enrichment score, cross-studies | maximize<br>maximize<br>maximize |
| Pre-clinical | RNASeq_LFC<br>RNASeq_pval | log2 fold change from internal RNAseq study, where cancer cell lines were treated with osimertinib vs DMSO treatment<br>adjusted p-values associated with LFC | maximize<br>minimize |

**Figure 6: SUPPLEMENTARY.**
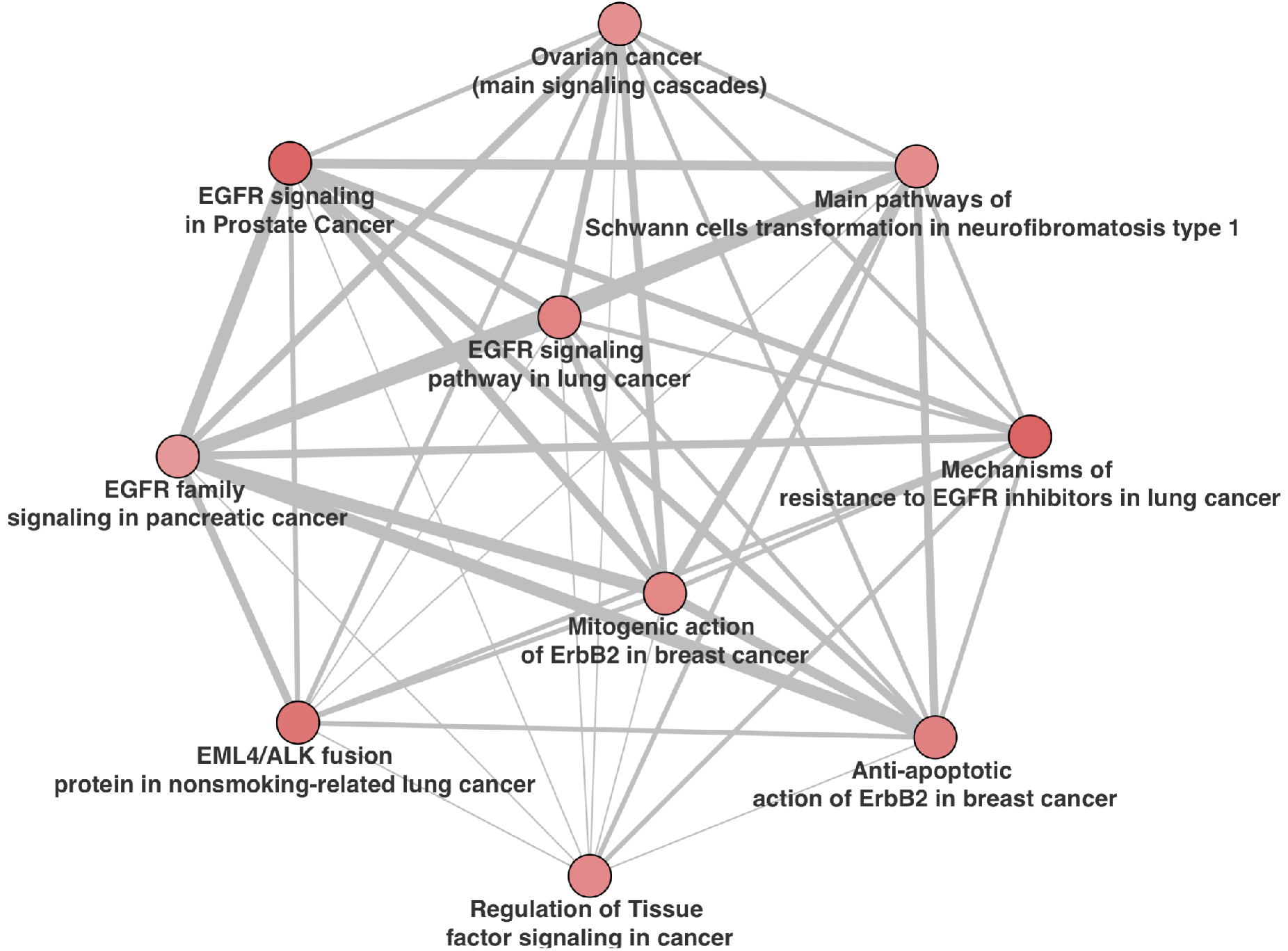
Crosstalk analysis of BIKG recommended hits: The network of the top 10 ontology terms from pathway enrichment analysis. Nodes represent top terms and edges represent significant similarities (as measured by hypergeometric test) between these entities. The edge thickness depends on the size of intersection between two ontology terms while the color of the node corresponds to the enrichment z-score.

**Table 2: SUPPLEMETARY.** Pathway Enrichment Analysis

| *Maps* | *Total* | *p-value* | *FDR* | *In Data* | *Network objects from data* |
| --- | --- | --- | --- | --- | --- |
| Mechanisms of resistance to EGFR inhibitors in lung cancer | 45 | 1.5e-13 | 1.6e-10 | 11 | ERK1 (MAPK3), ErbB2, K-RAS, PTEN, c-Src, ERK2 (MAPK1), AKT(PKB), E-cadherin, IGF-1 receptor, HGF receptor (Met), ErbB3 |
| Main pathways of Schwann cells transformation in neurofibromatosis type 1 | 80 | 1.2e-10 | 6.3e-08 | 11 | Tuberin, PDGF receptor, ErbB2, K-RAS, PTEN, ERK1/2, PDGF-R-beta, Neurofibromin, AKT(PKB), IGF-1 receptor, ErbB3 |
| EGFR signaling in Prostate Cancer | 46 | 2.4e-10 | 7.4e-08 | 9 | Tuberin, ERK1 (MAPK3), ErbB2, K-RAS, ERK1/2, c-Src, ERK2 (MAPK1), AKT(PKB), ErbB3 |
| Development_H3K36 demethylation in stem cell maintenance | 20 | 2.8e-10 | 7.4e-08 | 7 | CDK4, p15, Bcl-6, p53, EZH2, p14ARF, p16INK4 |
| Aberrant B-Raf signaling in melanoma progression | 55 | 1.3e-09 | 2.7e-07 | 9 | Tuberin, CBP, AKT1, ERK1/2, ROCK, ERK2 (MAPK1), BMF, AKT(PKB), Ric-tor |
| Regulation of Tissue factor signaling in cancer | 43 | 3.8e-09 | 6.6e-07 | 8 | ERK1 (MAPK3), K-RAS, VEGFR-2, PTEN, ERK1/2, ERK2 (MAPK1), AKT(PKB), p53 |
| Development_Estrogen-independent activation of ESR1 and ESR2 | 44 | 4.6e-09 | 6.9e-07 | 8 | ERK1 (MAPK3), ErbB2, CBP, ERK1/2, ERK2 (MAPK1), AKT(PKB), IGF-1 receptor, ErbB3 |
| LKB1 signaling pathway in lung cancer cells | 45 | 5.5e-09 | 7.3e-07 | 8 | Tuberin, PTEN, ERK1/2, c-Src, BRG1, p53, E-cadherin, p16INK4 |
| Main genetic and epigenetic alterations in lung cancer | 48 | 9.4e-09 | 1.1e-06 | 8 | K-RAS, CDK4, PTEN, p53, IGF-1 receptor, HGF receptor (Met), p14ARF, p16INK4 |
| The role of KEAP1/NRF2 pathway in skin sensitization | 32 | 1.1e-08 | 1.2e-06 | 7 | ERK1 (MAPK3), TXNRD1, CBP, ERK1/2, ERK2 (MAPK1), E-cadherin, KEAP1 |

**Figure 7: SUPPLEMENTARY.**
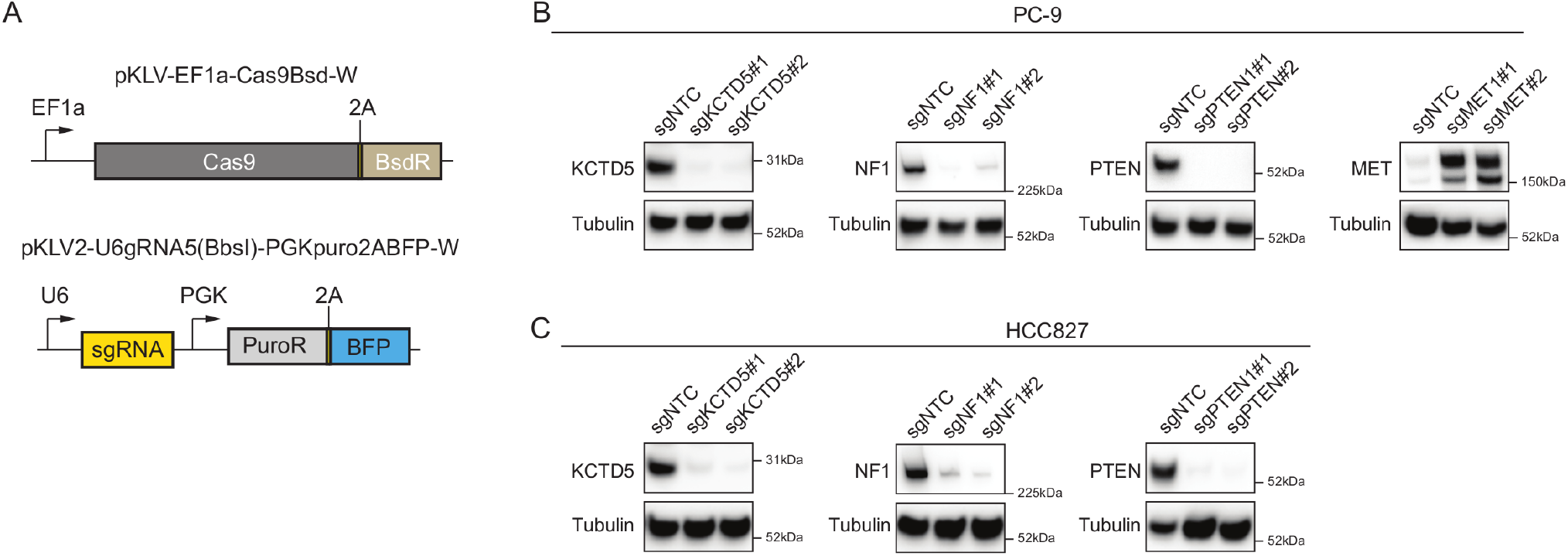
Evaluation of gene KO and activation in NSCLC cell lines. **A** CRIPSR Cas9 vector system used for generation of PC-9 and HCC827 KO cell lines. In these cell lines Cas9 is expressed constitutively after Blasticidin selection. pKLV2 sgRNA vector co-expressed BFP for FACS based tracking of proliferation. **B** Loss of target gene expression of KCTD5, NF1 and PTEN as well as overexpression of MET in PC-9. Whole cell lysates were analysed 14 days after transduction. **C** Loss of target gene expression of KCTD5, NF1 and PTEN in HCC827. Whole cell lysates were analysed 14 days after transduction.

**Figure 8: SUPPLEMENTARY.**
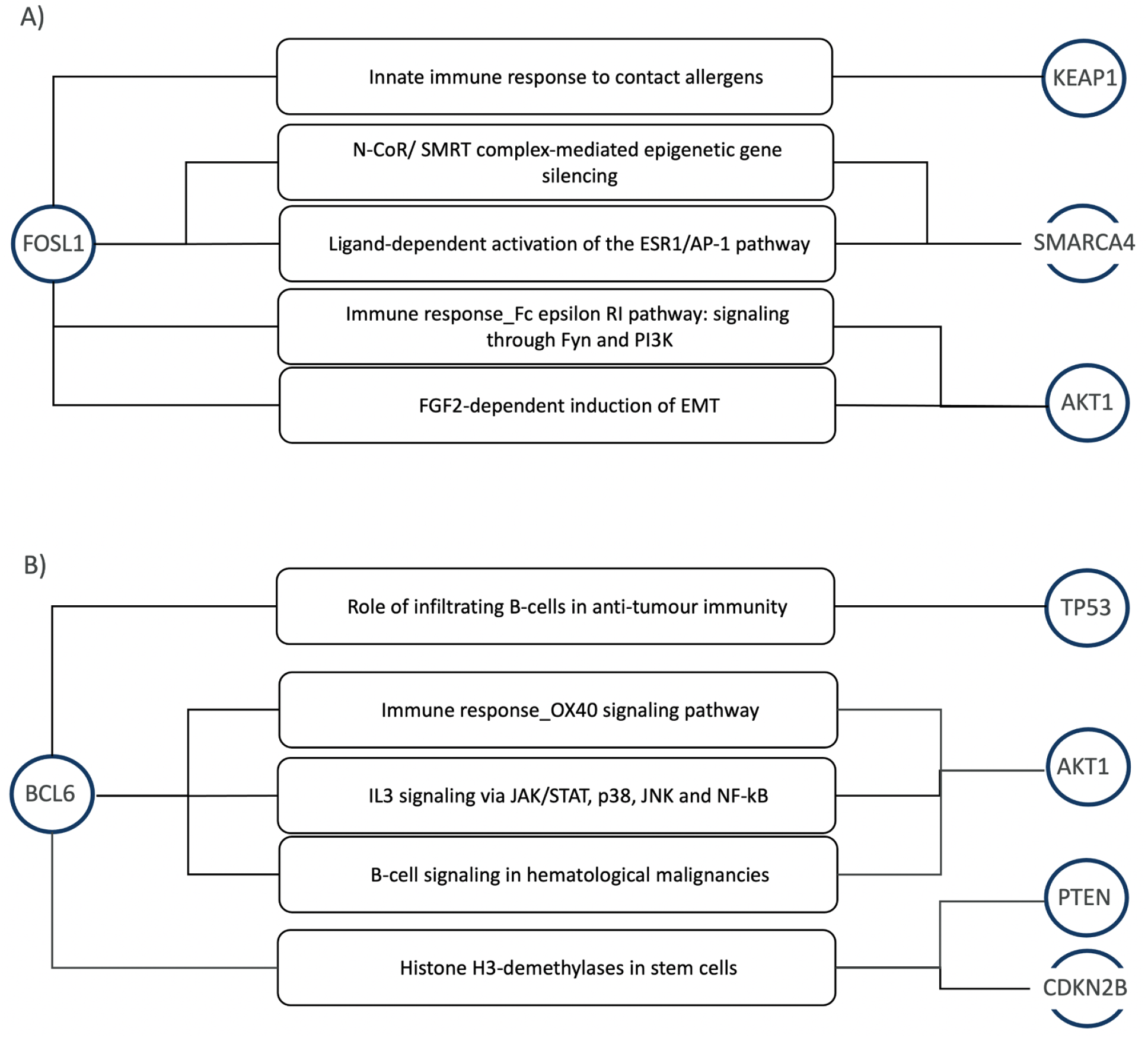
Novel markers predicted by recommendation system showing evidence paths to known osimertinib resistance markers. **A** shows FOSL1’s role in epigenetic silencing and innate immune response. **B** shows BCL6’s role in anti-tumour immunity, cross-talk with key aberrant pathways in cancer such as JAK/STAT and p38.

**Table 3: SUPPLEMENTARY.**
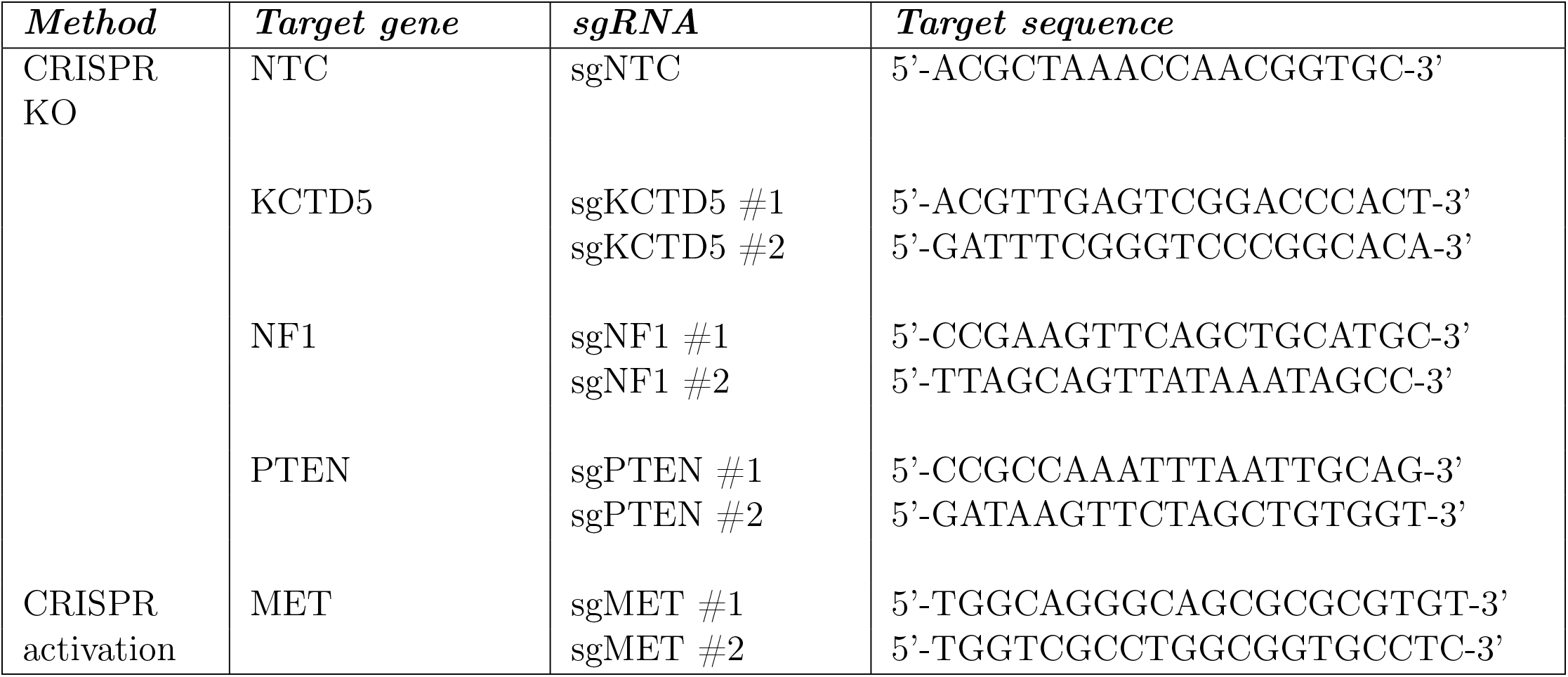
Target sequences of guide RNAs used in this study

